# Early life stress impairs postnatal oligodendrogenesis and adult emotional behaviour through activity-dependent mechanisms

**DOI:** 10.1101/369660

**Authors:** A. Teissier, C. Le Magueresse, J. Olusakin, B. L. S. Andrade da Costa, A. M. De Stasi, A. Bacci, Y. Kawasawa, V. A. Vaidya, P. Gaspar

**Affiliations:** INSERM, Institut du Fer à Moulin, U839 Sorbonne Université, Paris, France; Sorbonne Université, Paris, France; Institut Jacques Monod, CNRS UMR 7592, Université Paris Diderot, France; Physiology and Pharmacology Department, Federal University of Pernambuco, Recife, Brazil; Institut du Cerveau et de la Moelle épinière, CNRS UMR 7225 – Inserm U1127, Sorbonne Université, Paris, France; Departments of Pharmacology and Biochemistry and Molecular Biology, Institute for Personalized Medicine, Penn State University College of Medicine, Hershey, Pennsylvania, USA; Department of Biological Sciences, Tata Institute of Fundamental Research, Mumbai 400005, India

## Abstract

Exposure to stress during early life (infancy/childhood) has long-term effects on the structure and function of the prefrontal cortex (PFC) and increases the risk for adult depression and anxiety disorders. However, little is known about the molecular and cellular mechanisms of these effects. Here we focused on changes induced by chronic maternal separation during the first two weeks of postnatal life. Unbiased mRNA expression profiling in the medial PFC (mPFC) of maternally separated (MS) pups identified an increased expression of myelin-related genes and a decreased expression of immediate early genes. Oligodendrocyte lineage markers and birthdating experiments indicated a precocious oligodendrocyte differentiation in the mPFC at P15, leading to a depletion of the oligodendrocyte progenitor pool in MS adults. We tested the role of neuronal activity in oligodendrogenesis, using designed receptors exclusively activated by designed drugs (DREADDs) techniques. hM4Di or hM3Dq constructs were transfected into mPFC neurons using fast-acting AAV8 viruses. Reduction of mPFC neuron excitability during the first two postnatal weeks caused a premature differentiation of oligodendrocytes similar to the MS pups, while chemogenetic activation normalized it in the MS animals. Bidirectional manipulation of neuron excitability in the mPFC during the P2-P14 period had long lasting effects on adult emotional behaviours and on temporal object recognition: hM4Di mimicked MS effects, while hM3Dq prevented the pro-depressive effects and short term memory impairment of MS. Thus, our results identify neuronal activity as a critical target of early life stress and demonstrate its function in controlling both postnatal oligodendrogenesis and adult mPFC-related behaviors.

## INTRODUCTION

The environment a child encounters during early life plays a critical role in normal brain development. Thus, early life stress (ELS), which includes various forms of child abuse and neglect, has repeatedly been shown to increase the risk to develop psychiatric disorders (1–4). Current evidence that ELS alters developmental trajectories is essentially based on brain imaging studies in adult patients who experienced childhood adversity. These studies showed altered brain activity and myelin content, notably affecting the medial prefrontal cortex (mPFC), which constitutes a critical hub for emotional control and cognition (5–8). Since the mPFC has a protracted postnatal development, it is a likely target for environmental challenges occurring in childhood.

To identify the pathophysiological mechanisms leading to ELS phenotypes, in particular as regards mood and anxiety disorders, several preclinical models have been developed (9,10). The most common involves daily maternal separation of rodent pups during the early postnatal period, a time when pups normally spend most of their time huddled in the mother’s nest (9,11). Maternal separation (MS) during the first postnatal weeks induces a permanent impairment of emotional and cognitive behaviours, together with an altered physiology and myelination of the mPFC (12–15). Strikingly, these behavioural and structural defects closely reproduce the phenotypes of humans exposed to childhood trauma (10,16). Several of these alterations are already apparent at juvenile or adolescent stages (14,17–20) but the causal mechanisms remain poorly understood.

Here, we focused on cellular and molecular changes occurring in the mPFC at the outset of ELS. We started with an unbiased transcriptomic screen of the mPFC, comparing MS with standard facility raised (SFR) pups. We identified major changes in gene pathways associated with myelination and neuronal activity. Turning to cellular analyses, we documented a precocious differentiation of oligodendrocyte progenitor cells (OPC) in P15 MS pups, which coincided with an increase in myelin-related transcripts, and a depletion of the OPC pool in adults. Considering the important role of neuronal activity in adult myelination (21) we tested the contribution of altered mPFC neuronal activity during development on maternal separation phenotypes using designed receptors exclusively activated by designed drugs (DREADDs) (22). We found that local and transient decrease in neuronal activity from P2 to P14 is sufficient to induce premature OPC differentiation, while the transient increase in neuronal activity prevents it. Additionally, we demonstrate that some of the behavioural symptoms induced by ELS in adulthood are modulated by changes in mPFC neuronal activity during development.

## MATERIAL AND METHODS

### Animals

All experiments performed in mice were in compliance with the standard ethical guidelines (European Community Guidelines and French Agriculture and Forestry Ministry Guidelines for Handling Animals-decree 87849). All procedures have been approved by the ethical commity of the region Ile de France (Comité Darwin, agreement 09047.04). All dams were first-time pregnant BALB/cJ Rj mothers shipped from Janvier Labs ∼7 days before delivery. At arrival, pregnant females were group-housed by 2 and litter sizes were homogenized from 8 to 14 animals with both genders represented. After weaning at P21, all the mice were group-housed (4-5 per cage), and kept under standard laboratory conditions (22 ± 1°C, 60% relative humidity, 12-12 hrs light–dark cycle, food and water *ad libitum*) in ventilated racks. Males and females were similarly exposed to maternal separation protocol (MS pups) and/or viral injections, but only males were used for the experiments while females were sacrificed at weaning except for: (i) a first cohort of MS animals in which both genders were behaviourally tested and (ii) females receiving viral injection at P1 and sacrificed at P2, P4, P6, P8 and P10 for time course of viral expression or at P8 or P14 to test for CNO efficacy using Egr1 staining.

### Maternal Separation Protocol

Maternal separation of pups from their dams was conducted daily between 1:00 and 4:00 P.M., starting at postnatal day 2 (P2) and terminating at P14. Briefly, dams were first removed from the home cage and placed into a clean cage, then pups, were collected and placed together into another clean cage. The two cages were separated at some distance to avoid vocalized communications. After 180 minutes, pups and then dams were returned to their home cages. Control animals were standard facility-reared (SFR) and did not receive extra handling. Animals were sexed at P1 to allow random but homogeneous distribution of males, a first cage change occurred at P2 and then every week.

### RNA-sequencing

For the transcriptomic profiling, bilateral mPFCs from SFR (n=12) and MS (n=12) animals were dissected manually at P15 (Figure 1), fast frozen on dry ice and collected in 100 µL of RNALater® (Ambion, Life Technology). 3 unilateral mPFCs from different litters were pooled to generate 4 samples per group. Samples were shipped to Penn State Hershey Genome Sciences Facility (USA). Total RNA was extracted using mirVana kit (Thermo Fisher Scientific) and a bead mill homogenizer (Bullet Blender, Next Advance) together with an equivalent mass of stainless steel beads (Next Advance) were used to homogenize the tissue. The cDNA libraries were prepared using SureSelect Strand Specific RNA Library Preparation Kit (Agilent Technologies) and RNA and cDNA qualities were assessed using BioAnalyzer Kits (Agilent Technologies, RNA 6000 Nano and High Sensitivity DNA). The 12 libraries were loaded onto TruSeq SR v3 flow cells on an Illumina HiSeq 2500 and run for 50 cycles using a single-read recipe (TruSeq SBS Kit v3). The Illumina CASAVA pipeline Version 1.8 was used to extract de-multiplexed sequencing reads and FastQC (version 0.11.2) was used to validate the quality of the raw sequence data. Additional quality filtering was performed using FASTX-Toolkit with a quality score cut-off of 20. Next, alignment of the filtered reads to the mouse reference genome (mm10mm10) was done using Tophat (version 2.0.9) allowing 2 mismatches. Raw read counts were calculated using HTSeq-count (23). The “remove unwanted variation” (RUVg) strategy (“RUVseq” R package, v3.1) as described by Risso et al. (24) was used along with edgeR (25) to identify differentially expressed genes (DEG) between MS and SFR animals. First we normalized the raw read counts by selecting a set of “in-silico empirical” negative controls, i.e., 5000 least significantly DEG genes based on a first-pass DEG analysis performed prior to normalization. Normalized read counts were applied, using the negative binomial GLM approach implemented in edgeR, using the Likelihood Ratio Test method. Significantly DEG were defined to be those with q-value < 0.05 calculated by the Storey et al. method (26).

**Figure 1:**
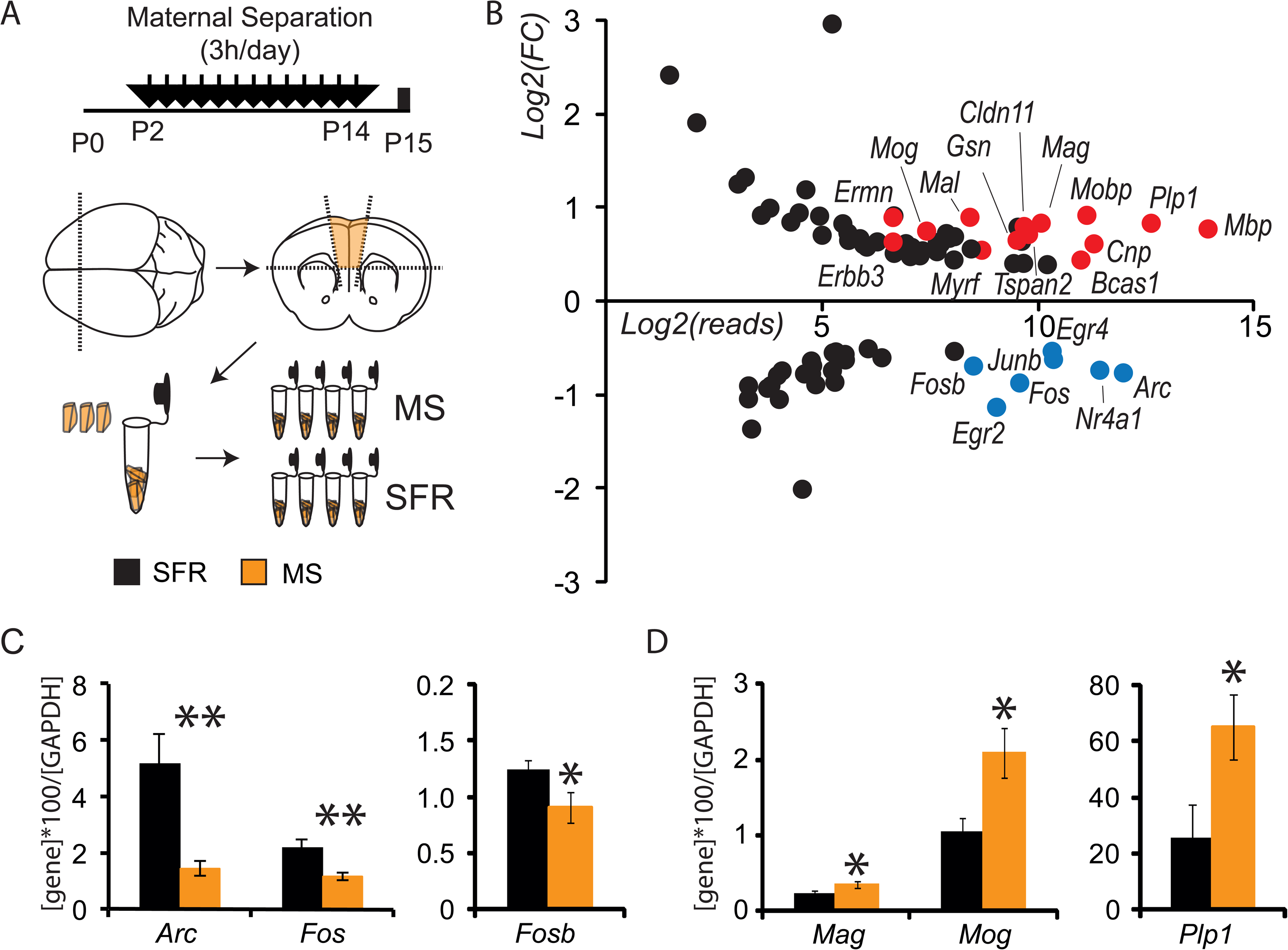
Analysis of differentially expressed genes in the mPFC of P15 animals exposed to maternal separation highlights myelin ensheathment and immediate early genes. (A) Schematic presentation of the experimental paradigm of the maternal separation (MS pups) versus standard facility (SFR pups) protocols and the dissection of mPFC at P15 to generate 4 samples per condition, each including 3 unilateral mPFC. (B) Graph shows with a log2 scale the fold change (FC) of expression induced by maternal separation according to the number of reads detected in control samples for the genes with q<0.05 and FC>1.3 or FC<0.7. Myelin-ensheathment genes are highlighted in red, and immediate early genes in blue. (C, D) Graphs show the qPCR results obtained in a second experimental cohort to determine the expression levels of IEG (C, n=7 SFR animals and 6 MS) and genes related to myelin ensheathment (D, n= 6 SFR animals and 9 MS). * p<0.05, ** p<0.01.

### RT-qPCR

mPFCs were dissected and homogenized in 1ml of TriReagent (Sigma-Aldrich) using a Polytron®PT13000D (Kinematica) and RNA was extracted according to standard protocols. Genomic DNA was removed by digestion with Amplification Grade DNase I (Sigma-Aldrich). First-strand cDNA was synthesized by reverse transcription of 5μg of total RNA with Superscript-II reverse transcriptase (Invitrogen) according to standard protocols. Reverse transcriptase was omitted in some samples as negative controls. Relative expression levels of *Gapdh, Arc, Fos, Fosb, Mag, Mog and Plp1* mRNAs were determined by real time quantitative PCR (RT-qPCR) using Absolute SYBR Green Mix (ABgene) and sets of specific primers (Supplementary Table 2) on a 96 wells plate (Thermo Scientific AB-0600). Gene expression was normalized to mouse *Gapdh* mRNA expression. Data were analysed with the 2–ΔCt method on MxPro qPCR software (Agilent Technologies), and values are expressed as the mean of duplicates.

### AAV injections in the PFC

AAV8-hSyn-hM3D(Gq)-mCherry, AAV8-hSyn-hM4D(Gi)-mCherry or AAV8-hSyn-GFP viruses (Addgene #50474, #44362 or #50465) were used as specified. The titres were within the range: 10^12^-10^13^ particles/ml. Injections were done on P1 mice pups anesthetized on ice (1minute or until the absence of pinch reaction) and maintained on a frozen pad during all the procedure (max 8 min). At the beginning of the experiment, litters were separated from the dams and placed into a heated cage with bedding. Pups were then individually anesthetized and maintained in a prone position with the skull flat, the skin was gently opened with a single incision with a razor blade. The mPFC was targeted using as a reference the inferior cerebral vein (∼2-3 mm posteriorly) and the superior sagittal sinus (∼2mm laterally). Bilateral injections (200 nl each) were performed 1.1 mm deep from the skull surface using a pulled glass capillary (30-50 µm tip diameter PCR micropipette, Drummond Scientific Company) mounted on a hydraulic micromanipulator MO-10 (Narashige, Japan). After the injection, the pups were immediately removed from the ice, the opened skin was sealed with tissue glue (Vetbond #1469, 3M, France) and pups were gently warmed up. All pups from the same litter were injected to avoid competition for feeding and were returned together to their home cage at the end of the procedure. Careful supervision was provided during the hours following the procedure to ensure mother acceptance and re-glue the scars if necessary. Afterwards, cages were returned to the normal facility and daily supervision was provided.

### Chronic CNO injections

A saline solution (SAL, 0.9% NaCl) or 5 mg/kg CNO + 0.9% NaCl solution was injected daily from P2 to P14, either subcutaneously (P2-P4) or ip (P5 onward). For SFR+hM4Di and SFR+GFP cohorts, injections were done at 1 pm. For the MS+hM4Dq cohort, injections were done at 4 pm at the end of the maternal separation period.

### Tissue processing

Mice were sacrificed at different postnatal times (P2-P15) and adult stages. All mice were anesthetized (Pentobarbital 0.5 mg/g) and fixed by intracardiac perfusion of 4% paraformaldehyde in 0.1 M phosphate-buffered (PB pH=7.2) using a peristaltic pump (Thermoscientific Masterflex). Brains were quickly removed, post-fixed overnight in the same fixative solution and cryoprotected 2 days in 30% sucrose containing 0.01% sodium azide. 35µm-thick coronal sections were prepared using a cryostat (Leica) or a cryotome (Microm Microtech). Serial sections from the whole brain were collected as series of 6. Tissue sections were used immediately for immunohistochemistry or stored at −20°C in a cryoprotective solution (30% ethylene glycol and 30% sucrose in 0.1 M phosphate buffer, pH=7.4).

### Immunohistochemistry

Immunolabelling techniques were used on free-floating sections, and all the washes and antibody incubations were performed in blocking solution containing 1% horse serum and 0.2% triton in PBS (PBST). The following primary antibodies were applied for 24 h (or 48 h for anti-cFos antibodies) at 4°C: mouse anti-APC/CC1 Ab-7 (Millipore, OP80, 1:100), chicken anti-mCherry (Abcam, 205402, 1:1000), rat anti-Ctip2 monoclonal (Abcam, ab18465, 1:500), rabbit anti-cFos (Synaptic System 226003, 1:1000) for P9 MS/SFR pups, rabbit anti-cFos antiserum (Abcam, 190289, 1:1000) for DREADDs experiments, mouse anti-NeuN (US Biological, 1:600), rabbit anti-Olig2 (Millipore Ab9610, 1:1000), rat anti-PDGFRα (BD Bioscience, 558774, 1:600) and rabbit anti-Tbr1 (Abcam, 1:1000). The corresponding fluorescent donkey antisera secondary (Jackson ImmunoResearch, 1:200) were applied for 2 h at room temperature. Sections were subsequently mounted using the SlowFade (Molecular Probes 536939).

### Image acquisition

Fluorescent images were acquired on a Leica SP5 confocal system equipped with an Argon laser (for the 488nm excitation), a Diode 561nm and HeNe 633nm. Z-series stacks of confocal images were acquired at 1024×1024 pixels resolution, with a pinhole set to one Airy unit and optimal settings for gain and offset.

### Western blot

P15 SFR and MS mice were decapitated and mPFC were macroscopically dissected on ice. Tissue from 2 brains were pooled and 600 μl of lysis buffer [Tris 100 mM pH 7.6, EDTA 0.5 M pH 8, 1% Triton X-100 and protease inhibitor cocktail (#P8340, Sigma, France)] was added to the tissue and sonicated on ice for 5 mins. Samples were centrifuged at 13,000 g (30 min at 4°C), supernatants were collected, and an aliquot from each sample was used for protein determination (bicinchoninic assay). A total of 40μg of each protein extract sample was subjected to SDS-PAGE, using NuPage 4–12% Bis-Tris polyacrylamide gels (Invitrogen, France). Proteins were blotted onto a 0.45-μm nitrocellulose membrane (Thermo Fisher, France). Membranes were blocked (1 h) in 5% non-fat milk dissolved in PBS (pH = 7.6) + 0.1% Tween (PBST) and incubated overnight in 1:1000 rabbit anti-PLP1 (Abcam, Cat. # ab28486) or 1:1000 mouse anti-Tuj1 (BioLegend, Cat. # 801201) at 4°C. After three rinses in PBST, membranes were incubated in Alexa Fluor 790-conjugated donkey anti-rabbit secondary antiserum (1:10000, Jackson ImmunoResearch, #711-655-152) or Alexa Fluor 680-conjugated donkey anti-mouse secondary antiserum (1:20000, Jackson ImmunoResearch, #715-625-151) at room temperature for 90 mins. Data were expressed as the ratio between the bands intensity of PLP1 and Tuj1.

### Acute slice preparation and electrophysiology

P9–P11 mice were sacrificed by decapitation and the brains were immediately removed from the skull and placed in oxygenated 4°C artificial cerebrospinal fluid (ACSF1), with the following concentrations in mM: 125 NaCl, 2.5 KCl, 25 glucose, 25 NaHCO_3_, 1.25 NaH_2_PO_4_, 2 CaCl_2_, and 1 MgCl_2_, and continuously bubbled with 95% O_2_-5% CO_2_. Slices (250 µm) were cut using a vibroslicer (HM650V, Microm) and incubated at 32°C for 20 minutes and then at room temperature (20-25°C). For patch-clamp recordings, slices were transferred to the recording chamber and continuously superfused with ACSF2 (30-32°C), with the following concentrations in mM: 125 NaCl, 3.5 KCl, 25 glucose, 25 NaHCO_3,_ 1.25 NaH_2_PO_4_, 1 CaCl_2_ and 0.5 MgCl_2_, pH 7.2 and continuously bubbled with 95% O_2_-5% CO_2_. mCherry^+^ neurons were visually identified using an upright microscope (BX 51, Olympus) equipped with standard epifluorescence. To minimize cell-to-cell variability, we selectively recorded from thick-tufted layer 5 pyramidal neurons which were identified based on their pyramid-like soma and stereotypical dendritic morphology with a prominent apical dendrite. Patch-clamp pipettes (4–6 Mohm resistance) were prepared from borosilicate glass (BF150-86-10; Harvard Apparatus) with a DMZ pipette puller (Zeitz). Patch-clamp experiments were performed using the following intracellular solution: 105 K-gluconate, 10 HEPES, 10 phosphocreatine-Na, 4 MgATP, 30 KCl (pH 7.25, adjusted with KOH). GABA_A_ receptors were blocked with 10μm SR95531 hydrobromide (Gabazine, HelloBio), AMPA receptors with 10μM 6-cyano-7-nitroquinoxaline-2,3-dione (CNQX, Hello Bio Inc), and NMDA receptors with 50 μM d-APV (HelloBio). To activate DREADDs in current-clamp experiments, mCherry^+^ thick-tufted layer 5 pyramidal neurons were patch-clamped in the presence of CNO (30 μM) or clozapine (1 μM) in the ACSF. Intrinsic excitability was measured as the firing frequency in response to incremental depolarizing current steps. To measure CNO-induced currents in voltage-clamp experiments, holding current was recorded every 2s. 30 consecutive sweeps starting 2 minutes after CNO application were averaged and compared to the average of 30 sweeps immediately prior to CNO application (“baseline”). Barium chloride (100 μM, Sigma) was always applied in the ACSF at the end of the experiment to block CNO-induced currents. The junction potential (−5mV) was not corrected for. Data acquisition was performed using Patchmaster software (Heka Elektronik). Signals were sampled at 20 kHz and filtered at 3 kHz, and off-line analysis was performed using Igor (WaveMetrics Inc.). The experimenter was blind to both treatment and/or viral subtype.

### Behavioural Studies

Female and male mice were tested separately for emotional behaviours starting at P80 and animals were generated to reach at least 10 per group. The tests were performed in the following order: open-field (OF), elevated plus maze (EPM), marble burying, sequential object recognition (SNOR), splash test and the two-days forced swim test, with 3 to 7 days between each test according to the stressful character of the test. All behavioural testing was done during the light cycle between 9 am and 6 pm. Animals were habituated to the testing room for at least 30 min and the experimenter was blind to animal treatment. To eliminate odour cues, each apparatus was thoroughly cleaned with alcohol after each animal. Testing took place under dim light conditions (40lux) for OF, EPM, SNOR and splash test to enhance tracking of white mice, and under normal light conditions (80 lux) for the other tests.

#### Open Field

mice were placed in the centre of rectangular plastic grey boxes (H24,5xL35xL55cm) and activity was recorded for 20 min. Total distance and time spent in the centre of the OF were automatically recorded using AnyMaze tracking software (Stoelting Co).

#### Elevated Plus Maze

the maze is a grey plus-crossed-shaped apparatus, with two open arms (30 cm) and two closed arms (30 m) of 5cm width, linked by a central platform and located 40cm above the floor. Mice were individually put in the centre of the maze facing an open arm and were allowed to explore the maze for 10 min. The time spent and the entries into the open arms were automatically measured using AnyMaze tracking software and used as anxiety indexes.

#### Marble Burying

housing cages (20×36 cm) were filled with 6 cm of clean wood bedding with 12 marbles uniformly spaced (27). Mice were individually placed in a corner of the cage and the number of marbles buried on more than 2/3 of their surface was manually recorded every minute for 20 min.

#### Splash Test

group housed animals were placed in a new cage with bedding for at least 30 min. Mice were then individually sprayed on the back twice (∼2×0,6 ml) with 20% sucrose solution in water and placed in a corner of their home cage. Grooming behaviours, including licking, stroking and scratching were manually recorded using AnyMaze software keys.

#### Sequential Novel Object Recognition

this is an adapted version of the novel object recognition test considered to be more specific to mPFC functioning (28). Animals were first habituated to the arena (same grey boxes as OF), without stimuli for 10 min daily for 2 d before the behavioural testing. The task comprised two sample phases and one test trial. In each sample phase, mice were allowed to explore two copies of an identical object for a total of 4 min. Different objects were used for sample phases 1 and 2 (miniature car then plastic hook, or glass tube then cork of Falcon tube, or plastic mug then cell culture flask), with a 1 h delay between the sample phases. The test trial (3 min) was given 3 h after sample phase 2 and consisted in placing together one copy of each set of objects. If temporal order memory is intact, the subjects will spend more time exploring the less recent object (from sample 1) compared with the more recent object (from sample 2). The positions of the two objects were counterbalanced between animals.

#### Porsolt Forced Swim test

Testing of behavioural despair was carried out on two consecutive days using a glass cylinder (40 cm×20 cm diameter) filled with water (23-25°C) up to 3/4. Mice were tested once a day for 6 min. All the swim sessions were videotaped and manually tracked using AnyMaze software keys. Only day 2 data are represented.

### Statistics

The Gaussian distribution of the values was tested on Prism software using d’Agostino and Pearson or Shapiro-Wilk for n<7. If data followed a Gaussian distribution, 2 tailed T-test or one-way ANOVA test were performed; otherwise specific tests were applied as specified in the text. All results are expressed in mean ± sem. * p<0.05, ** p<0.01 and *** p<0.005.

## RESULTS

### Early life stress (ELS) alters the expression of immediate early genes and genes involved in myelin formation in the developing medial prefrontal cortex (mPFC)

As a preclinical model of ELS, we used the common model of chronic maternal separation, where pups are daily separated for 3h from their dam starting at postnatal day 2 (P2) and until P14 (Figure 1A) (11,15). This procedure has previously been shown to cause behavioural phenotypes, such as increased depressive-like symptoms and cognitive deficits, suggestive of alterations of mPFC development. To identify early molecular candidates potentially involved in the permanent alterations induced by ELS, we focused on the chronic effects of maternal separation by analysing P15 brains, sampled 24h after the last maternal separation (MS), compared with standard facility raised (SFR) pups. We performed an unbiased genetic screen using high-throughput RNA sequencing and minimized inter-individual variations and dissection biases by pooling mPFC of 3 males from different litters and testing 4 independent samples per condition. After normalization, we found a total of 86 genes differentially expressed between MS and SFR animals (adjusted p-value <0.05 and change in transcript level >30%, Supplementary Table 1). Among the 56 up-regulated genes, Gene Ontology (GO) analyses revealed a major enrichment in biological processes related to myelin ensheathment and oligodendrocyte differentiation (red dots in Figure 1B and Supplementary Figure 1A). Similar analyses of the 30 down-regulated genes showed enrichment in cellular response to calcium (Supplementary Figure 1B) likely due to a strong representation of several immediate early genes (IEG) (blue dots in Figure 1B). To validate these changes we performed quantitative PCRs on mPFC mRNAs obtained from a new cohort of MS and SFR animals analysed at P15. This second cohort confirmed the decrease in of several IEG transcripts (*Fos*, *Fosb* and *Arc*, Figure 1C) and the increase in the myelin-related transcripts analysed (*Mag*, *Mog* and *Plp1*, Figure 1D). We further examined whether increase in gene expression was reflected at protein levels. Western blots against PLP1 in the P15 mPFC showed an increase in PLP1 levels after MS (0,45 +/-0,07; n=5) compared to SFR (0,27 +/-0,06; n=5; p<0,01) (Supplementary Figure 1C,D). Altogether, this unbiased approach identified an early concomitant alteration of transcripts related to IEG and myelin ensheathment in the mPFC of postnatal animals exposed to maternal separation.

### Premature OPC differentiation in ELS pups

To determine possible causes of the increase in myelin-related transcripts, we investigated oligodendrogenesis in the MS animals. The postnatal period corresponds to an intense proliferation of oligodendrocyte progenitor cells (OPCs) and P15 marks the initiation of differentiation into myelinating oligodendrocytes in the mPFC (20,29). We performed triple immunostaining of Olig2, PDGFRα, and CC1 to distinguish OPCs (Olig2^+^PDGFRα^+^, Figure 2A) from differentiated oligodendrocytes (OLs, Olig2^+^CC1^+^) (30). At P15, the density of OPCs was unaltered in the mPFC of MS animals compared to SFR animals (F(1,7)=0.032 p=0.863, Figure 2B), but we detected a significant increase in the density of OL (F(1,7)=21.84 p=0.0023 Figure 2C). At adult stages, we observed a significant decrease in OPC density (F(1,8)=6.28 p=0.036 Figure 2D), while no difference in OLs density or Plp1 levels was detected after MS (F(1,8)=0.022 p=0.8865, Figure 2E and Supplementary Figure1D). To test whether early differentiation contributes to these observations, sequential staining of the dividing progenitors was performed using two thymidine analogues at P10 (iodo-deoxyuridine, IdU) and P15 (bromo-deoxyuridine, BrdU, Figure 2F). This interval was chosen because OPCs differentiate into OLs in approximately 5 days (29). In the mPFC of P15 MS animals, the fraction of OPCs that kept dividing between P10 and P15 (triple labelled with Olig2^+^IdU^+^BrdU^+^, blue dots in Figure 2G, F(1,5)=12.62 p=0.0163) was significantly reduced, while the differentiating fraction (Olig2^+^IdU^+^BrdU^−^, pink dots, F(1,5)=7.234 p=0.0433) was increased. Thus, our results demonstrate that maternal separation induces a precocious differentiation of the OPC pool during development, as a potential cause for depletion of the OPC pool in adults. To assess how oligodendrogenesis defects could be related with the alterations in IEG we previously observed, we performed immunostainings on MS P9 mPFC (Supplementary Figure 2). We observed that the decrease in c-Fos expression was already present at P9, while no change in CC1^+^Olig2^+^ or PDGFRα^+^Olig2^+^ density could yet be detected. These results suggest that the chronic down-regulation of neuronal activity in the MS pups preceded the defects in oligodendrogenesis. Considering the important role of neuronal activity in adult myelination (21), we further tested how manipulation of neuronal activity over the early postnatal period could modify oligodendrogenesis.

**Figure 2:**
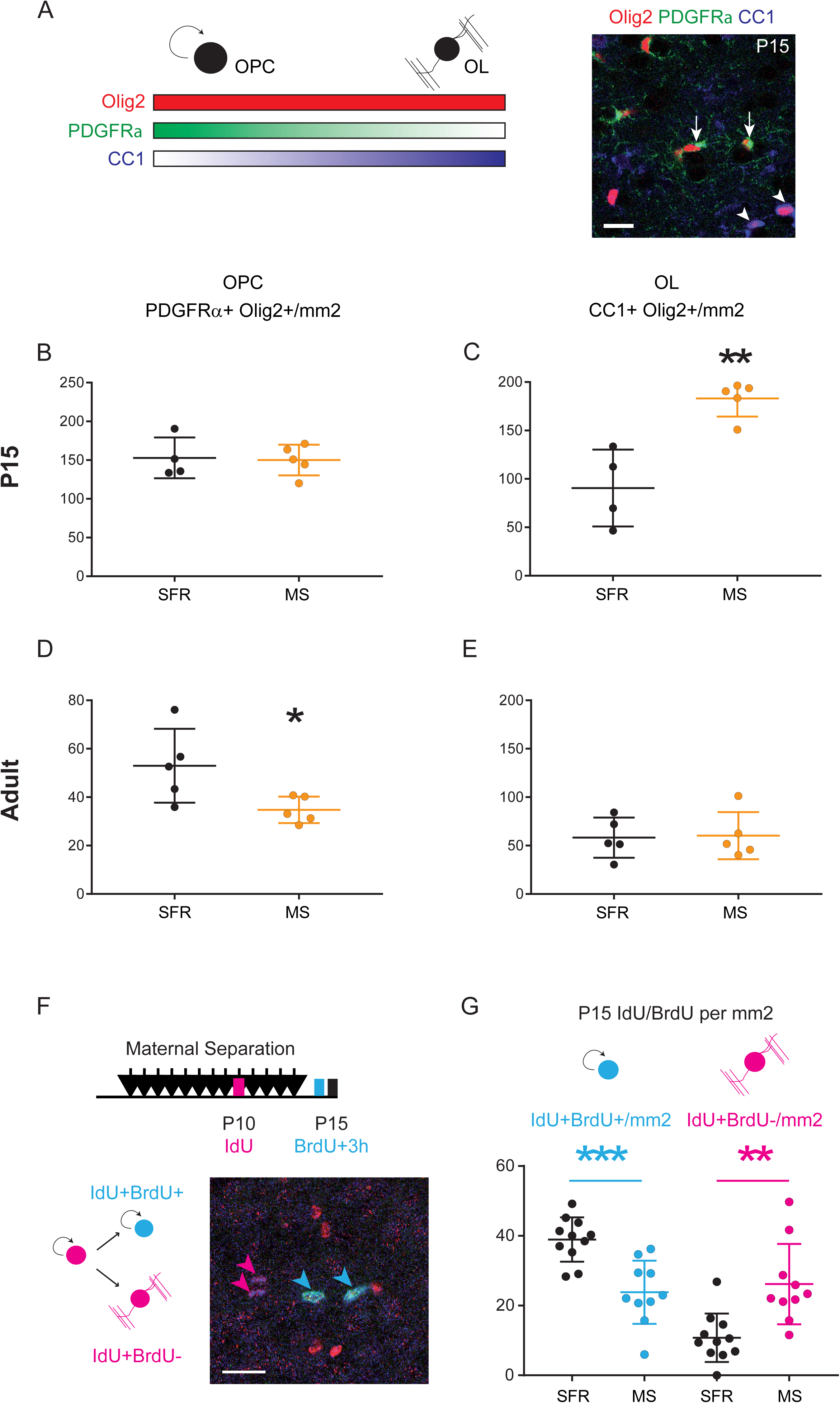
Maternal separation induces precocious differentiation of oligodendrocytes in the developing mPFC. (A) The scheme illustrates the specificity of the oligodendrocyte lineage markers using triple immunostaining. Oligodendrocyte progenitor cells (OPCs) express Olig2 and high levels of PDGFRα while mature oligodendrocytes (OLs) express Olig2 and high levels of the CC1 antigen. Scale bar represents 20µm. (B, C, D, E) Graphs show the density of OPCs (B,D) and OLs (C,E) in the P15 (B,C n=5 animals per conditions) and adult (D,E n=5 animals per conditions) mPFCs of SFR and MS animals. (F) Scheme shows the protocol of sequential labelling of OPCs using IdU injections at P10 and BrdU injections at P15 to differentiate the fraction of P10 OPCs that are still cycling at P15 (blue, IdU^+^BrdU^+^) from those which have started differentiation by then (purple, IdU^+^BrdU^−^). (G) Graph shows an increased fraction of differentiated OPCs in P15 mPFC of MS animals as compared to SFR animals (n= 11 SFR animals and 10 MS animals). * p<0.05, ** p<0.01, *** p< 0.005.

### Chemogenetic manipulation of mPFC postnatal activity bi-directionally controls developmental oligodendrogenesis

We used viral delivery of AAVs expressing DREADDs to transiently and bi-directionally manipulate neuronal activity in the mPFC (22,31). AAV8-hSyn-hM4Di-mCherry or AAV8-hSyn-hM3Dq-mCherry viral constructs were bilaterally injected into the mPFC of all P1 pups (Figure 3A). A time course analysis of DREADD expression in the mPFC showed mCherry expression visible as early as P2, with a progressive increase until P6 when expression reached a plateau (Figure 3B). The characterization of cell types transfected was done at P15: 38%±5.2 of the mPFC NeuN^+^ cells were mCherry^+^ upon hM4Di injections and 51.8%±4.4 upon hM3Dq injections. In addition, 95,98%±0.06 of the mCherry^+^ cells were NeuN^+^, and no co-expression of mCherry with Olig2 or PDGFRα or the oligodendrocyte*-*specific antibody CC1 could be detected (data not shown) supporting that our viral transfection experiments essentially targeted neurons. A bias for deep-layer glutamatergic neurons was observed using Tbr1 and Ctip2^+^ as markers of these populations: 72,59%±3.75 of the mCherry^+^ cells were Tbr1^+^ or Ctip2^+^ and only 6.17%±1.24 were GABA^+^ (Figure 3B-C).

**Figure 3:**
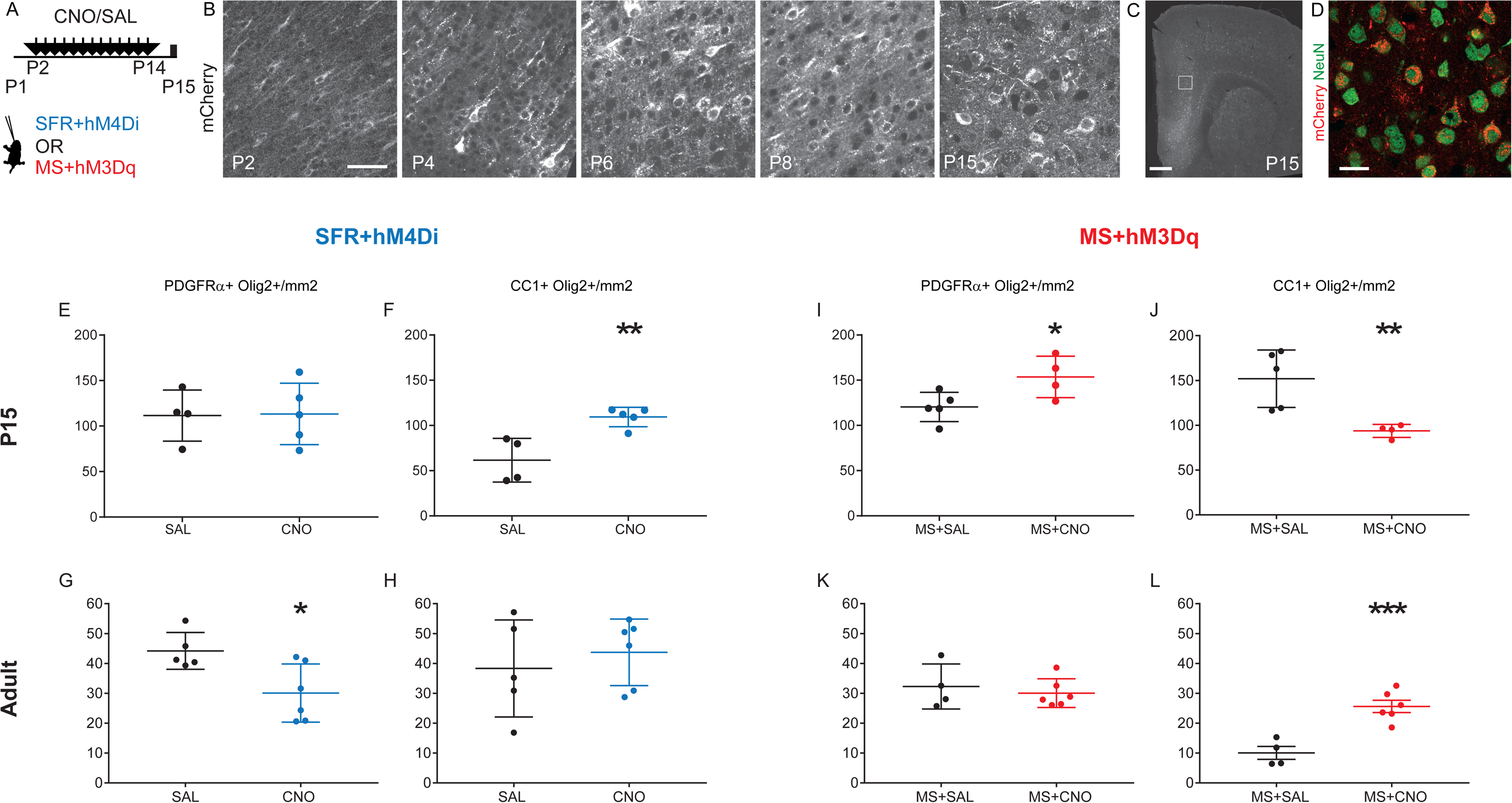
Neuronal activity bi-directionally controls oligodendrocytes differentiation in the developing mPFC. (A) Scheme illustrates the experimental paradigms of the chronic treatment with CNO or SAL solutions from P12 to P14 after local and bilateral injections of viral constructs expressing DREADDs and fused mCherry. All animals were either injected with the inhibitory hM4D(Gi) virus and raised under standard facility protocol (blue, SFR+hM4Di), or injected with excitatory hM3D(Gq) virus and maternally separated (red, MS+hM3Dq). (B) Time course of DREADDs expression in the postnatal mPFC as observed by mCherry labeling. (C, D) Micrographs show that transfection of the constructs in brain section of P15 mPFC (B) was restricted to neuronal lineage as indicated NeuN and mCherry co-labelling (C). (E-H) Graphs show the density of OPCs (E, G) and OLs (F, H) at P15 (E, F, n=4SAL/5CNO) and in adults (G, H, n= 5SAL/6CNO) mPFCs of SFR+hM4Di. (I-L) Graphs show the density of OPCs (I, K) and OLs (J, L) at P15 (I, J n=5SAL/4CNO) and in adults (K, L n= 4SAL/6CNO) mPFCs of MS+hM3Dq animals. Scale bars represent 50µm, (B), 500 µm (C) and 20 µm (D).* p<0.05, ** p<0.01, *** p< 0.005.

All mice received viral injections in the PFC at P0, and further received daily injections of saline (SAL) or the inert ligand Clozapine-N-Oxyde (CNO) from P2 to P14. Pups injected with the inhibitory construct were raised in the SFR condition (referred to as SFR+hM4Di, Figure 3A). Conversely, pups injected with the activating construct were submitted to the maternal separation protocol (referred to as MS+hM3Dq).

To evaluate how these experimental conditions modified PFC activity, we used *ex vivo* patch-clamp electrophysiological recordings to measure neuronal excitability and immunolabeling of immediate early genes (c-Fos/Egr1) to evaluate the *in vivo* consequences on mPFC neural activity. We first tested the effect of DREADD activation on intrinsic excitability, i.e. the propensity of a neuron to fire action potentials when subjected to an input current. In slices of animals receiving mPFC hM3Dq injections, there was a robust increase in neuronal intrinsic excitability in the hM3Dq-mCherry^+^ layer 5 pyramidal neurons upon exposure to CNO (30µM) or to its potential metabolite clozapine (1µM) (Supplementary Figure 3A,B). In parallel, a significant increase in cFos labelling in the mPFC was observed (Supplementary Figure 3I). In slices prepared from pups with the SFR+hM4Di protocol, exposure to CNO or clozapine did not modify intrinsic excitability significantly (Supplementary Figure 3C,D), although the low firing frequency and the resulting high trial-to-trial variability in our experimental conditions may have made small effects difficult to detect. Since hM4Di activates G protein-coupled inwardly-rectifying potassium (GIRK) channels, we tested whether CNO elicits GIRK-mediated currents in voltage-clamped hM4Di-mCherry^+^ pyramidal neurons. Indeed CNO application (30 μM) elicited a small but significant outward current (16.3 ± 5.8 pA, n=11, p=0.002, Mann-Whitney test) that was inhibited by barium (100 μM), a blocker of GIRK channels (Supplementary Figure 3E,F). The inhibitory action of hM4Di was also confirmed by a significant decrease in mPFC Egr1 expression among hM4Di-mCherry^+^ neurons in P8 and P15 animals sacrificed 2 hours after CNO injection (Supplementary Figure 3G). Additionally, a sustained decrease in cFos labeling was observed in P15 mPFC of SFR+hM4Di mice sacrificed 24h after chronic CNO exposure (P2-P14) compared to the saline treated group (Supplementary Figure 3H). Altogether, these results support the efficacy of our approach for either mimicking or counter-acting the chronic reduction in IEG expression in the mPFC induced by MS during the postnatal period.

To test the role of changes in early neuronal activity in the oligodendrogenesis defects induced by maternal separation, P15 and adults brains were analysed after SFR+hM4Di or MS+hM3Dq treatments, as described above. In P15 SFR+hM4Di animals exposed to CNO, we found no difference in OPC density but an increased density of OLs compared to saline exposed animals (F(1,7)=15.991 p=0.005, Figure 3F); while at adult stages we observed a decrease in OPC density (F(1,9)=7.809 p=0.0209, Figure 3G) with no changes in OL density in CNO/SAL groups. Conversely, in the P15 MS+hM3Dq mPFC, we found an increase in OPC density (F(1,7)=6.52 p=0.0379, Figure 3I), with no changes in OL density in CNO/saline group comparisons (Figure 3J). In adults that had been developmentally exposed to MS+hM3Dq there was an increase in OL density (F(1,8)=25.795 p=0.001, Figure 3L) with no alterations in OPC density when comparing the CNO/SAL groups (Figure 3K). Overall, this indicated that transient reduction of mPFC neuron excitability reproduced the oligodendrogenesis defects of MS mice, suggesting a similar premature exit of the cell cycle. Moreover, increasing neuronal excitability of mPFC neurons was sufficient to prevent the oligodendrogenesis defects induced by MS. These results indicated a causal role of altered mPFC neuronal activity on the precocious oligodendrogenesis observed in MS animals.

### Postnatal local decrease in mPFC activity recapitulates MS behavioural phenotype in adults

We then tested whether chemogenetic manipulations in the developing PFC could affect the emotional and cognitive symptoms induced by MS. We began with a characterization of the behavioural phenotype of adult MS BALBcJ mice (Figure 4A) since some variability of long term behavioural effects of MS have been reported between different mouse strains (11). In our protocol, adult MS males showed decreased exploration (Mann-Whitney p=0.0002) and time spent in the centre in the open field (OF, F(1,25)=4.317 p=0.0482, Figure 4B and C) as well as decreased time spent in the open arm of the elevated plus maze (EPM, F(1,22)=5.438 p=0.0293, Figure 4D). They further showed an increased burying behaviour in the marble test (F(1,32)=5.402 p=0.0266, Figure 4E), when compared with SFR animals. In addition, MS males displayed an increased time floating in the forced swim test (FST, F(1,26)=6.048 p=0.0209, Figure 4F) and an impaired short-term memory in the sequential novel object recognition test (SNOR, F(1,23)=4.75 p=0.0396,) which measures temporal order memory (as the preference in exploring an object seen less recently than another) (18,28). These results confirmed an increase in anxiety and depression-related behaviours in MS animals and an impairment in a test of recognition memory, which has been linked to mPFC function (28).

**Figure 4:**
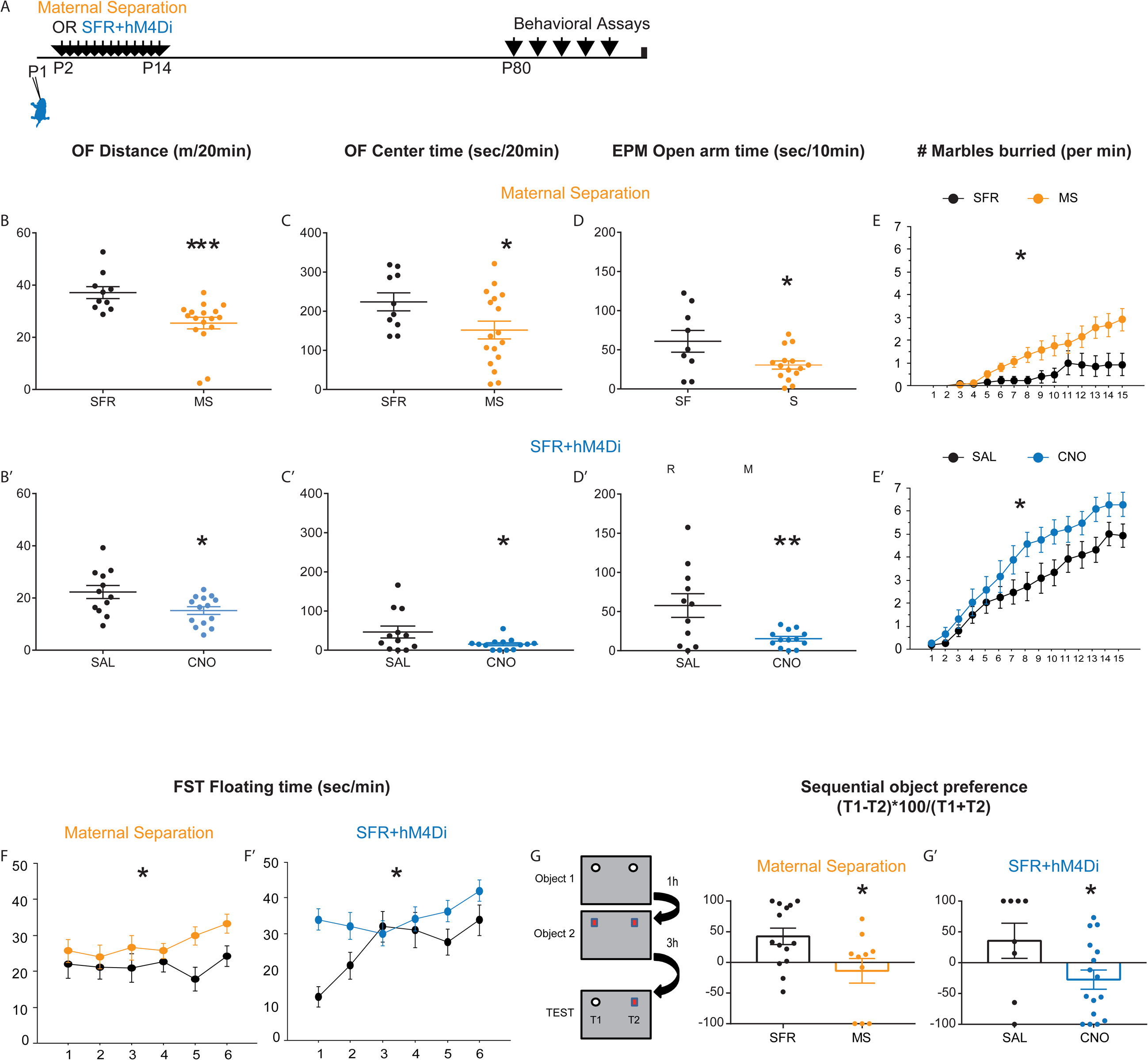
Early inhibition of mPFC neurons recapitulates the emotional disorders induced by early life stress. (A) Scheme illustrates the experimental paradigms of maternal separation (orange) or SFR+hM4Di (blue) treatments followed by behavioural testing in adulthood. (B,B’) Graphs show the distance travelled in an open field (OF) by SFR or MS animals (B, n=10SFR/17MS) and by SFR+hM4Di animals exposed to chronic SAL or CNO (B’, n=12SAL/14CNO). (C, C’) Graphs show the time spent in the center of the OF during the same 20min test. (D, D’) Graphs show the time spent in the open arm of an Elevated Plus Maze (EPM) during a 10min test (n= 9SFR/15MS animals and n=11SAL/13CNO). (E, E’) Graphs show the number of buried marbles at each minutes of a 15min test (n=14SFR/20MS and n=13SAL/15CNO animals). (F, F’) Graphs show the time floating per min of a 6 min forced swim test (FST, n=10SFR/18MS and n=12SAL/16CNO). (G, G’) Scheme represents the experimental paradigm of the sequential object recognition test with 3min exposure to two identical object1, followed 1h later by 3min exposure to two identical object 2, and 3h later by 4min exposure to object1 and object2. Graphs indicate the preference ratio for Object1 (n=14SFR/10MS animals and n=8SAL/15CNO animals). * p<0.05, ** p<0.01, *** p< 0.005.

In adults SFR+hM4Di animals, comparison of mice exposed to CNO or SAL from P2 to P14 revealed behavioural alterations very similar to those of MS animals. These included reduced exploration in the OF (distance F(1,24)=6.433 p=0.0181, Figure 4B’ and centre time F(1,24)=4.46 p=0.0452, Figure 4C’), reduced exploration in the EPM (F(1,22)=9.94 p=0.0067, Figure 4D’), increased marble burying (F(1,26)=5.233 p=0.305, Figure 4E’), increased immobility in the FST (F(1,26)=5.675 p=0.0248, Figure 4F’) and decreased preference in the SNOR test (F(1,19)=4.443 p=0.0485, Figure 4G’). They also show delayed postnatal growth with a sustained decrease in adult weight, as MS animals (Supplementary Figure 4). To exclude any unwarranted effects of the effects of chronic exposure to CNO, we injected a new cohort of animals with AAV8-hSyn-GFP at P1, administered CNO or SAL daily from P2 to P14 and tested their adult behaviour. In this cohort, no change in any of the behavioural tests was found between the CNO and SAL groups (Supplementary Figure 5). Altogether, our data show that decreasing mPFC neuronal activity during development is sufficient to recapitulate salient features of the MS’s adult phenotype.

### Postnatal local increase in mPFC activity prevents the emergence of depression-related phenotype induced by maternal separation

To test the role of reduced mPFC activity in the behavioural phenotype of MS animals, we assessed the behaviour of animals exposed to MS with or without chemogenetic activation (hM3Dq) of mPFC neurons (Figure 5A). CNO and SAL treated MS mice showed no difference in the behavioural parameters related to anxiety including: the distance or the time in the centre of the OF (Figure 5B, F(1,32)=0.604 p=0.4427 and C, F(1,32)=1.397 p=0.246), the time in the open arm of the EPM (Figure 5D, F(1,33)=2.002 p=0.1664) or the number of burying in the marble test (Figure 5E, F(1, 32)=1.656 p=0.2074). However, MS+hM3Dq animals exposed to CNO floated less in the FST (Figure 5F, F(1,29)=6.508 p=0.0163) and had improved performance in the temporal recognition memory-test (Figure 5G, Mann-Whitney p=0.0463). To strengthen our analysis of the depression-related phenotype we used the sucrose splash test, which measures self-care and hedonic behaviours (32). Animals solely exposed to MS display a decrease in grooming when compared with SFR animals (Figure 5I, F(1,25)=5.618 p=0.0258). We observed an increased grooming behaviour in CNO exposed MS+hM3Dq animals as compared to SAL (Figure 5H, F(1, 25)=6.532 p=0.171). Hence, transient increase in mPFC neuronal excitability prevented the emergence of the depression and recognition memory defects that characterize MS exposed mice, but had no effect on the anxiety-related phenotypes.

**Figure 5:**
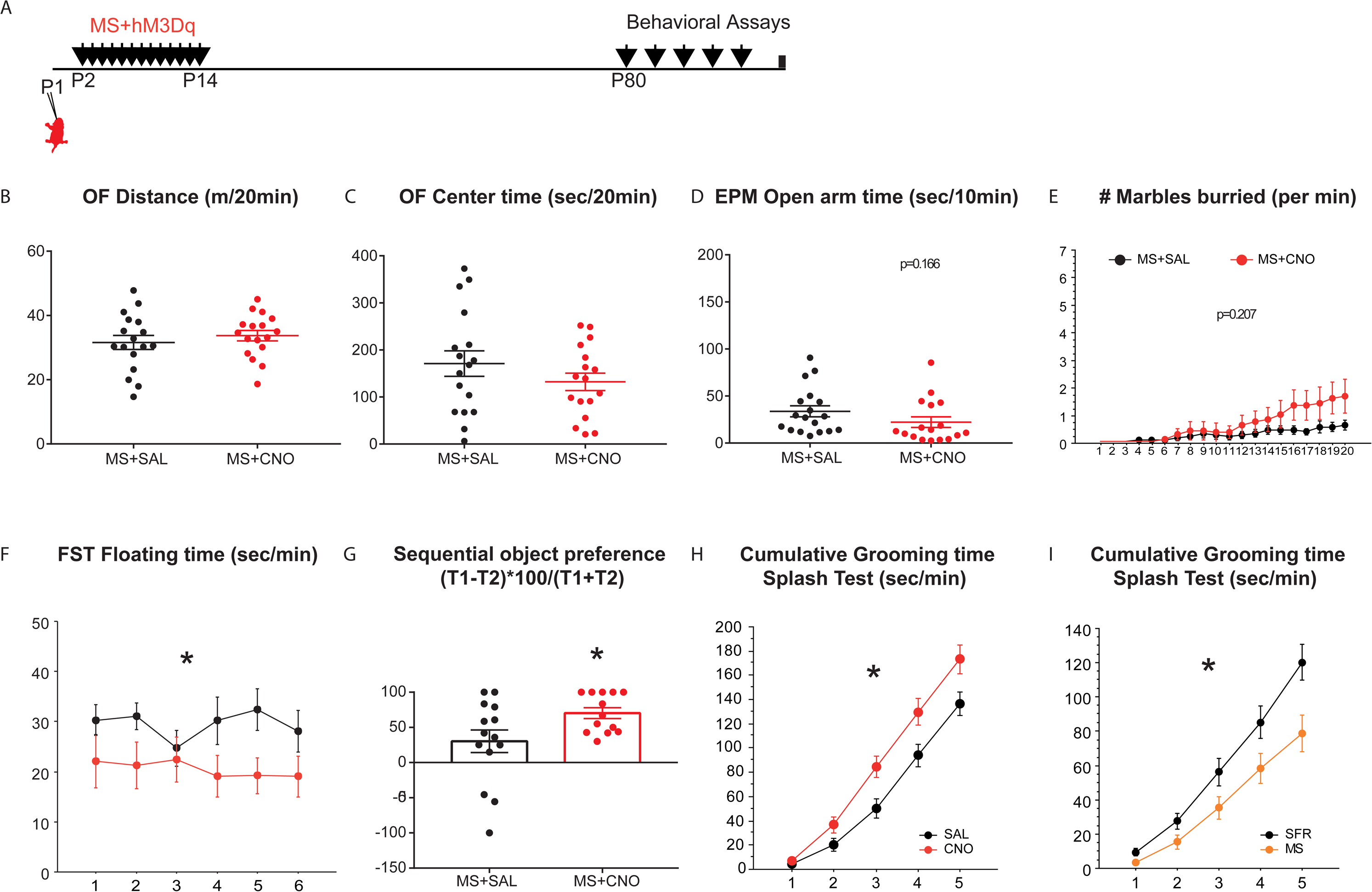
Early activation of mPFC neurons prevents the emergence of the depression-like phenotype induced by early life stress. (A) Scheme illustrates the experimental paradigms of MS+hM3Dq animals, analysed for behaviour as adults. (B) Graph shows the distance travelled in an open field (OF) by MS+hM3Dq animals exposed to SAL (black, n=17) or CNO (red, n=17). (C) Graph shows the time spent in the center of the OF during the same 20min test (n=17SAL/17CNO). (D) Graph shows the time spent in the open arm of an EPM during a 10min test (n=18SAL/17CNO). (E) Graph shows the number of buried marbles at each minutes of a 20min test (n=18SAL/16CNO). (F) Graph shows the time floating per min of a 6 min FST. (G) Graph shows the preference ratio for Object1 (n=14SAL/13CNO). (H, I) Graphs show the cumulative time grooming after a splash with 20% sucrose solution. Comparison was made between SFR+hM3Dq animals exposed to chronic SAL or CNO (H, n=14SAL/13CNO) and between SFR or MS animals (I, n=15/12). * p<0.05.

## DISCUSSION

The present study identified alterations in the expression of genes related to neuronal activity and to myelination in the developing mPFC of mice exposed to early life stress. We found that chronic maternal separation induced a precocious differentiation of the oligodendrocyte lineage in pups, together with a subsequent depletion of the OPC pool in adults. Chemogenetic tools demonstrated that these alterations are controlled by local changes in mPFC neuronal activity during early postnatal development. Furthermore, we established that bidirectional changes of neuronal activity in the developing mPFC are sufficient to modulate adult behaviours, and in addition, to mimic or rescue the long-term effects of early life stress.

### The pre-weaning period is critical for mPFC myelination

Early life stress in humans has been linked with hypo-myelination in the adult prefrontal cortex (5–8), a phenotype also reported in rodent models of ELS(9–12,20). Present findings provide a possible developmental mechanism for this, by showing that a precocious differentiation of the oligodendrocytic lineage during development could be at the origin of the permanent reduction in the adult OPC pool. Thus, the first 2 postnatal weeks appear to represent a critical period for oligodendrogenesis, with an activity-dependent regulation of maturation and cell cycle exit that has long term effects on the adult progenitor pool. This could add up to later critical periods for oligodendrogenesis identified in juveniles, where social isolation (P21-P35) also induced a decrease in myelin content and immediate early gene’s expression which correlate with a permanent impairment in social behaviour at adult stages, although the developmental mechanisms were unclear (33). Importantly, developmental exposure to stress contrasts with the effects of adult exposure to chronic stress which has only transient consequences on mPFC myelination and behaviour (33–35). Therefore, our results, by showing the importance of the P2-P14 period, suggest that successive and/or expanded critical period(s) occur in the mPFC to establish mPFC myelination, and correlate with alterations in the functions of anxiety, mood and social circuits.

The exact developmental mechanisms involved in adult hypomyelination after ELS are still disputed (7,12). Our data favour a precocious differentiation of the oligodendrocyte lineage resulting in a depletion of the OPC pool in adults. In another study, Yang et al. (20) reported a deficit in OPC differentiation in the mPFC of MS Sprague Dawley rats. The discrepancy with the present results is unclear and could be due to a number of technical differences between the studies such as delayed timing of analysis (P15 in mice vs P21 in rats) and the differences in markers of proliferation that were used. Importantly, our findings are supported by a transient increase in PLP1 levels as well as by several others, previous observations in other experimental models of sensory deprivation or ELS, showing transient and immediate increase in myelin content, followed by a permanent decrease in adulthood in sensory cortices or cerebellum (36–40). Moreover, a recent clinical paper indicating that childhood adversity in human biases oligodendrocyte lineage toward a more differentiated phenotype in the ventral mPFC, with more pronounced effects in samples from young individuals that died closer to ELS experience (7). Altogether, these results suggest that early stressors induce a pathological precocious differentiation of OL, which thereafter alters myelin content, or the capacity of myelin plasticity in adult life.

### Neuronal activity controls early stages of oligodendrogenesis

Our unbiased approach identified an important decrease of several immediate early genes in the P15 mPFC after maternal separation, as a proxy for a deficit in neuronal plasticity (41). The converging reduction of *Fos* and *FosB* further suggest that there is a chronic dampening of neural network activity in the mPFC of pups undergoing chronic maternal separation experience. These results are in agreement with previous reports in the MS rat model at P21, showing a local decrease in specific immediate early gene expressions correlating with a reduced neuronal synchrony in the mPFC, as revealed by *in vivo* recordings (14,18). Converging experimental evidence, provided in the present study, support a causal link between activity changes and the precocious differentiation of oligodendrocytes. On one hand, reduced cFos expression in the mPFC preceded the changes in OPCs differentiation (Supplementary Figure 2), on the other hand, chemogenetic inhibition of the mPFC in the early postnatal period induced a bias toward OPC differentiation, thus mimicking the MS phenotype. Finally, increasing neuronal excitability prevented the bias toward OPC differentiation, induced by MS. To the best of our knowledge, this is the first demonstration of a causal role of neuronal activity on mPFC oligodendrogenesis during the postnatal life.

In adults, increased neuronal activity stimulates proliferation and differentiation of OPCs and increases myelination in motor cortices (21,42,43), while chronic stress decreases IEG expression and impairs OPC proliferation and myelination in the mPFC (33,44). This is opposite to the effect that we observe during early postnatal life, since we show that increased neuronal activity stimulates proliferation at the expense of differentiation. Therefore, it is likely that different activity-dependent regulations of oligodendrogenesis are at play during development. This hypothesis is also supported by changes in the molecular properties of OPCs during this postnatal period (45,46), including the establishment of transient synaptic connectivity between OPCs and interneurons (21,47). It will be important in the future to determine how activity-dependent signalling differs during early postnatal life and adulthood to control OPC maturation and which are the cellular substrates involved.

### Early mPFC physiology and psychiatric disorders

Another important finding of the present study is the direct role of postnatal mPFC activity in the aetiology of emotional and memory phenotypes induced by ELS.. The mPFC neuronal network is still being assembled during the postnatal period, and previous studies have identified P2-P14 as a critical period for emotional and cognitive functions (48). A critical role of neuronal activity in cortical maturation is well established in the primary sensory cortical maps (visual, somato-sensory, auditory) (49,50). Activity thus appears as the mechanistic link between changes in the environment and alterations in the brain’s developmental trajectory. Our study provides the first evidence of an activity-dependent maturation of mPFC emotional circuits. An important question remains however open, which is determining the effects of ELS on neural circuit activity, which is presently unclear, although some recent investigations have started addressing this question with *in vivo* physiological recordings. Using this technique in P12-P17 rats, these studies showed that ELS decreased the power of low frequencies oscillations in the mPFC and altered cortical synchrony(51,52). These effects could be driven by changes in neuronal activity in different mPFC circuits, as suggested by another recent study showing that optogenetic activation of deep layers generates a global increase in mPFC oscillatory power in the neonatal mouse cortex (P7), while activation of the upper layers mainly drives beta oscillations (53). These recent technical advances raise hope for future dissection of ELS pathophysiology and could help resolving whether altered maturation of specific mPFC circuits could underlie vulnerability to depression and anxiety-disorders

Our results also suggest that developmental mechanisms underlying anxiety- and depression-related behaviors differ, as mPFC activation in MS pups only restored the depression-related phenotype. The reason for this is unclear. Generally one may argue that restoring a normal phenotype requires more precision in the activation of spatial and temporal activity patterns, whereas inhibition is more crude. In addition, our viral infection showed a bias toward deep layers containing PFC-raphe connections important in depression-related phenotypes (54), whereas the PFC-amygdala connectivity sustaining anxiety originate from both superficial and deep mPFC layers (55).

On a translational perspective, our results demonstrate that early mPFC activity tightly controls two major endophenotypes related to ELS, namely oligodendrogenesis and depression-related behaviour, thus supporting a common origin for these typical traits of emotional disorders (56–58). Strikingly, most mental disorders, including emotional disorders and schizophrenia, involve alterations of mPFC myelination and the susceptibility to develop these disorders is increased by ELS (56,59,60). The possible causal role of oligodendrogenesis in the aetiology of emotional disorders has been suggested by several recent studies. For instance, genetic alterations of oligodendrogenesis were correlated with emotional disorders in adult rodents (33,61–65) and restoring myelination is sufficient to rescue the social avoidance induced by chronic stress (35). Present results further support this notion, by showing that ELS alters mPFC oligodendrogenesis and emotional behaviours via shared activity-dependent mechanisms.

## Conclusion

This study demonstrates that ELS leads to a dampening of immediate early gene expression in the developing mPFC, resulting in a precocious maturation of the oligodendrocyte lineage and a subsequent reduction of adult myelin progenitors in adult life. Moreover, our results show that early postnatal changes in mPFC neuronal activity in mouse pups play a crucial role in determining the emotional and temporal memory alterations that emerge in adulthood after early life stress events.

## Contributions

A.T. generated the cohorts, performed the histological and behavioural experiments and analysed all the data. J.O. and B.L.S.A.d.C. provided help for the preparation of animal cohorts and histological analysis. C.L.-M. and A.d.S. performed the electrophysiological recordings and analysed the data. Y.L. performed and analysed the transcriptome profiling experiments. A.B. and V.A.V. helped to design the study. P.G. and A.T. designed the study and wrote the manuscript with the help of all authors.

## Acknowledgements

We thank Dr. Macklin, Dr. Soiza-Reilly, Dr. Parras and Dr. Angulo for their scientific and technical advices and comments on the manuscript. We are also thankful to the members of the CEF UMS28 facility as well as the U839 and U894 imaging platforms for their support.

## Funding and disclosure

The work was funded by the Labex BioPsy, the French Agence Nationale pour la Recherche (ANR grant FRONTELS), the DIM Région Ile de France, and the Behavior and Brain Foundation Young Investigator-NARSAD Grant. B.A.d.C. received a scholarship from Coordination for the Improvement of Higher Education Personnel (CAPES), Brazil. J.O. is the recipient of the Ecole des Neurosciences de Paris (ENP) scholarship.

The authors have no conflict of interest to report.

## Notes

#### Summary of Updates

revised version of the manuscript after reviews from Molecular Psychiatry

