## Supplementary Figures for "Early life stress impairs postnatal oligodendrogenesis and adult emotional behaviour through activity-dependent mechanisms"

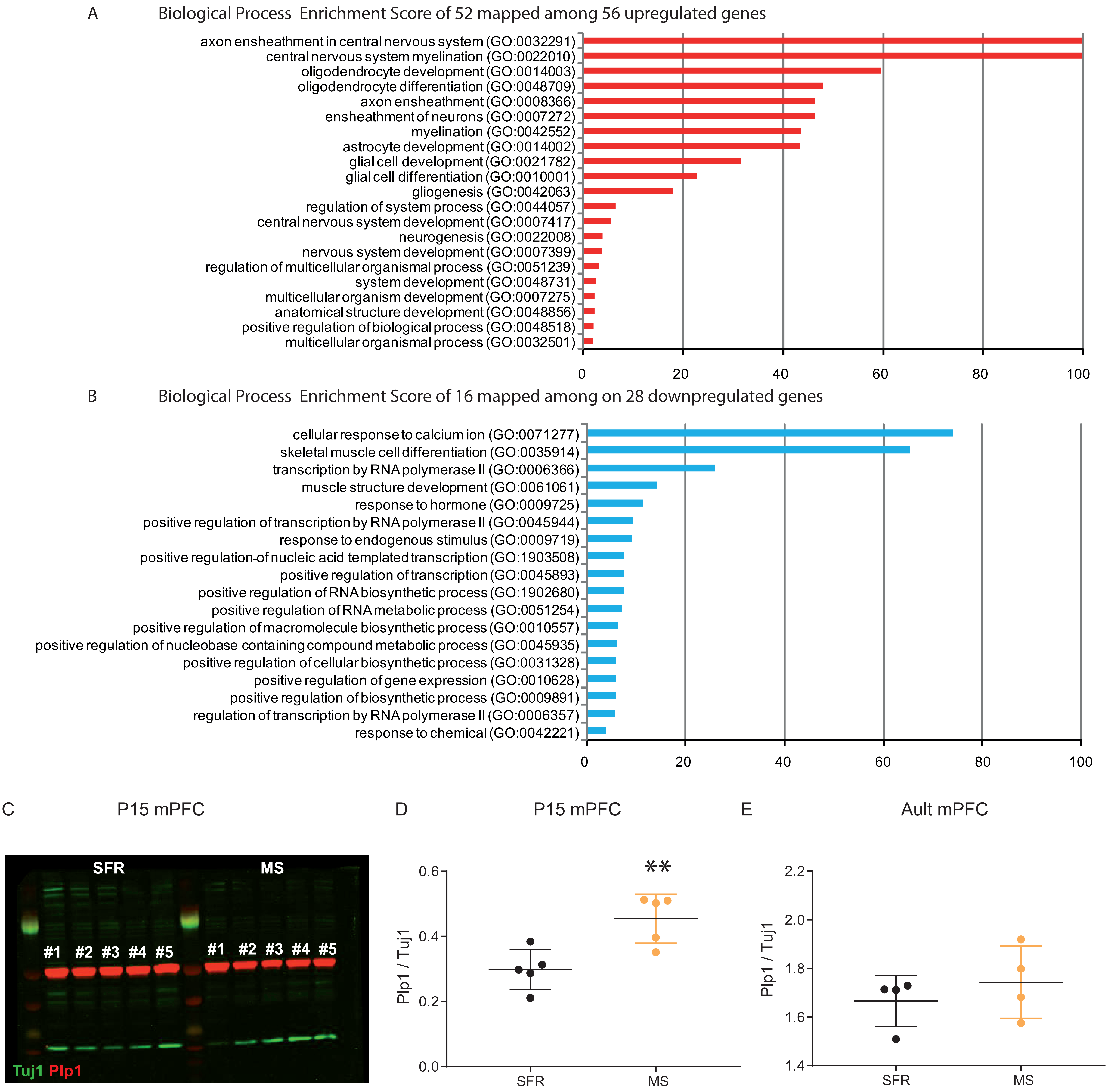


**Supplementary Figure 1**

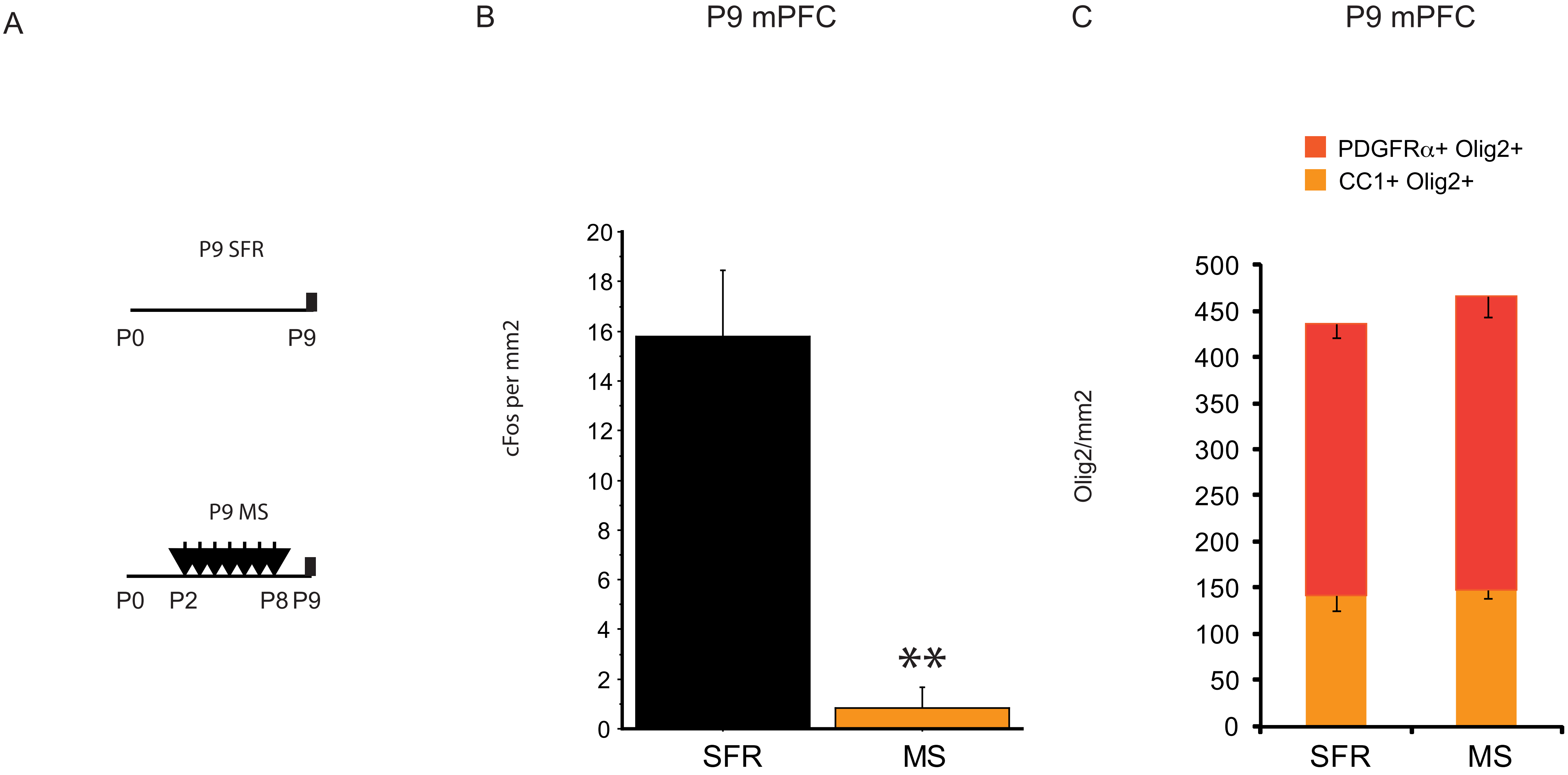


. **Supplementary Figure 2**

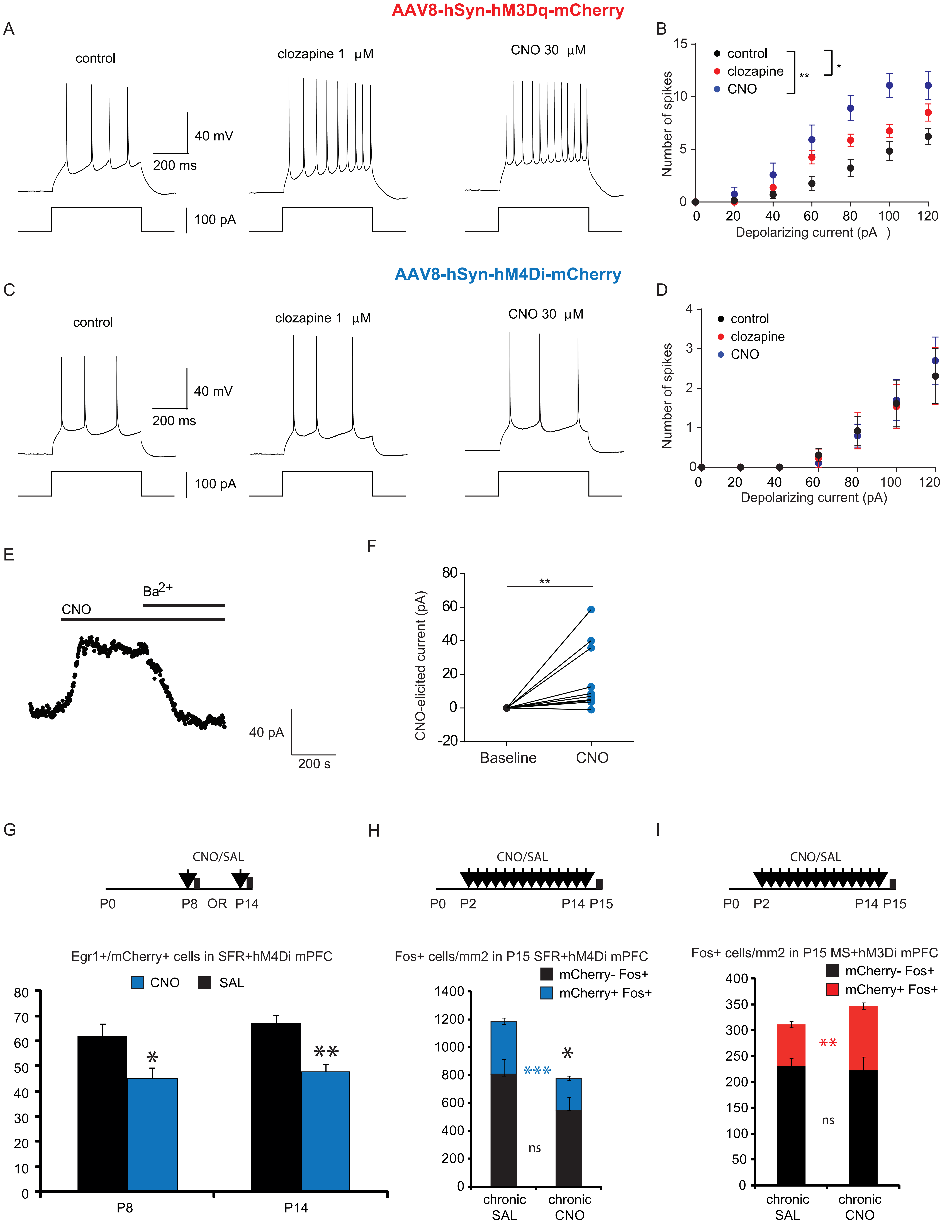


**Supplementary Figure 3**

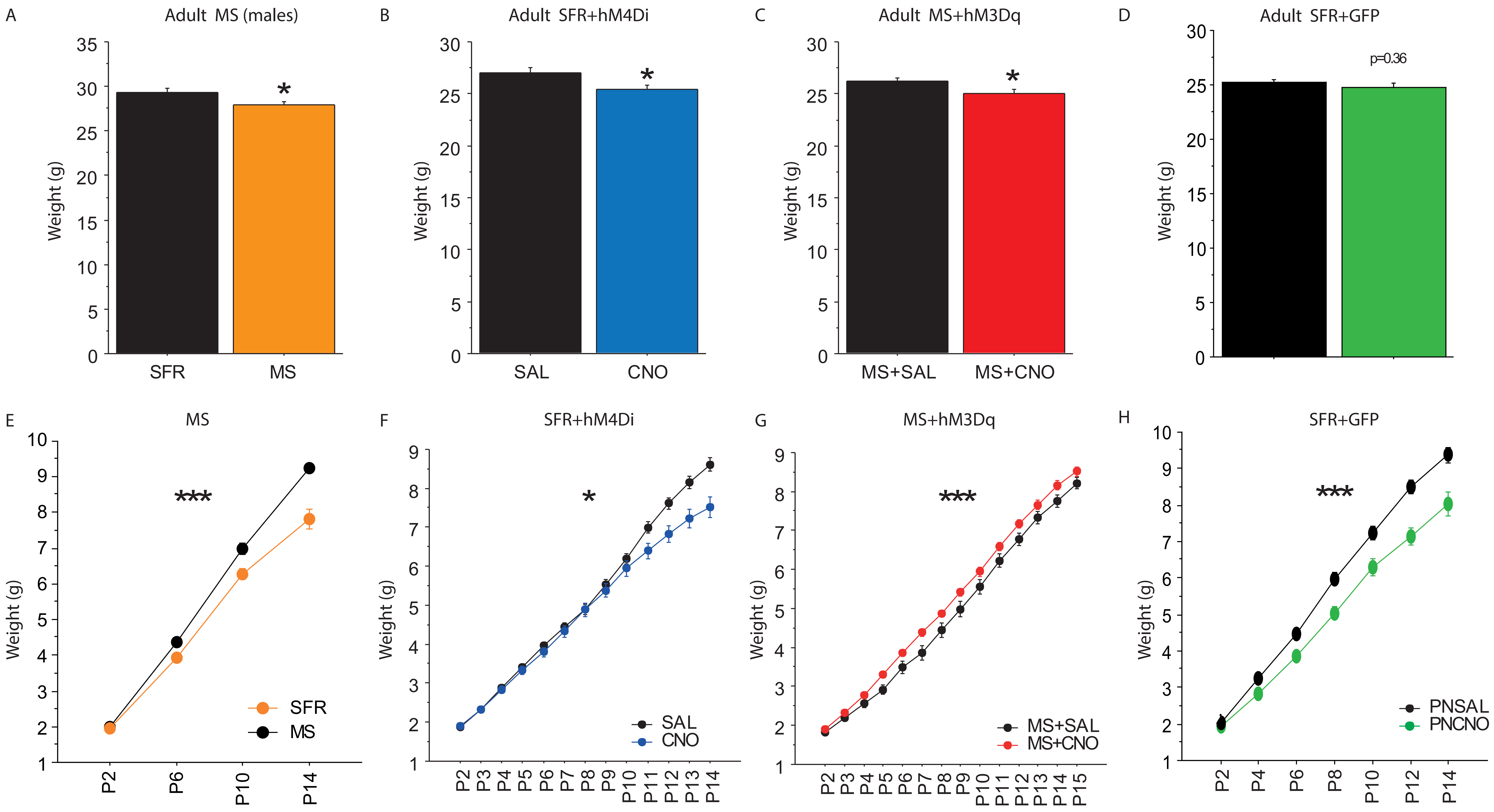


**Supplementary Figure 4**

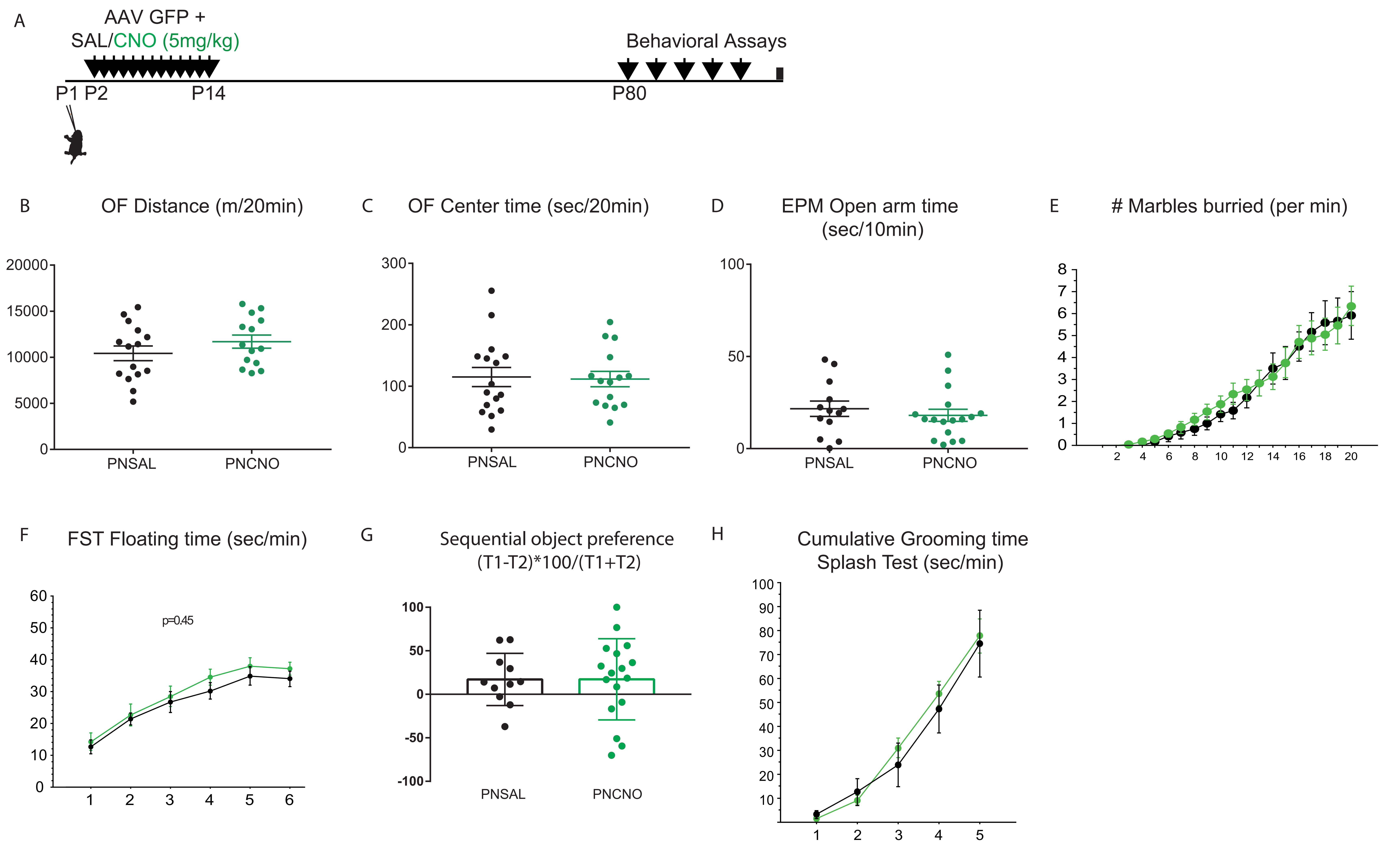


**Supplementary Figure 5**

### Supplementary Table 1: Differentially Expressed Genes after MS in P15 mPFC (q<0.05)

| Associated Gene Name | LOG2 reads SFR | LOG2 FC MS | p value | qvalue |
|---|---|---|---|---|
| Egr2 | 9,02375383 | -1,126006315 | 6,5E-21 | 1,08E-16 |
| Nr4a1 | 11,41156395 | -0,730383731 | 2,78E-17 | 2,3E-13 |
| Tspan2 | 9,762062828 | 0,720010208 | 5,77E-12 | 2,39E-08 |
| Ugt8a | 9,517559001 | 0,797271877 | 1,1E-11 | 3,64E-08 |
| Scn4b | 6,648903197 | 0,918592049 | 1,57E-11 | 4,35E-08 |
| Egr4 | 10,30941143 | -0,528635473 | 1,11E-10 | 0,000000264 |
| Junb | 10,33996104 | -0,615224159 | 2,17E-10 | 0,00000045 |
| Gjc3 | 7,652885845 | 0,632713474 | 2,52E-10 | 0,000000464 |
| Cd4 | 1,460527981 | 2,42098154 | 3,26E-10 | 0,000000541 |
| Fos | 9,558588375 | -0,866091025 | 7,89E-09 | 0,00000872 |
| Ier2 | 8,048671807 | -0,529595203 | 8,93E-09 | 0,00000925 |
| Gm28437 | 5,221248586 | 2,967628059 | 9,59E-09 | 0,00000936 |
| Arc | 11,95602984 | -0,758643317 | 0,000000011 | 0,0000101 |
| St18 | 4,613254194 | 1,195836658 | 1,21E-08 | 0,0000106 |
| Lpar1 | 7,933875798 | 0,678589785 | 2,45E-08 | 0,0000203 |
| Prlr | 4,532633011 | -2,001629386 | 0,000000213 | 0,000139417 |
| Plp1 | 12,60011609 | 0,83878217 | 0,000000219 | 0,000139417 |
| Ermn | 6,62903443 | 0,900631705 | 0,000000396 | 0,000234541 |
| Nts | 4,842267279 | -0,885197351 | 0,000000442 | 0,000252899 |
| Myrf | 8,681915138 | 0,551147447 | 0,000000568 | 0,00030364 |
| Mag | 10,05480175 | 0,840152772 | 0,000000562 | 0,00030364 |
| Fosb | 8,49062432 | -0,685752069 | 0,000000633 | 0,000327936 |
| Eomes | 2,087515124 | 1,912468164 | 0,000000867 | 0,000435788 |
| Gm13066 | 5,526470294 | -0,620020444 | 0,00000102 | 0,000487098 |
| Mobp | 11,11124947 | 0,923463751 | 0,00000103 | 0,000487098 |
| Adamts4 | 7,72845064 | 0,620668354 | 0,00000139 | 0,000604872 |
| Plxnb3 | 8,43739156 | 0,56462054 | 0,00000235 | 0,000950779 |
| Nr1i3 | 6,054142033 | -0,500907386 | 0,00000305 | 0,001204231 |
| Clic6 | 3,997992816 | -1,042206393 | 0,00000355 | 0,00133789 |
| Mog | 7,405370509 | 0,755453762 | 0,00000348 | 0,00133789 |
| Ppp1r1b | 9,604652043 | 0,646620602 | 0,00000391 | 0,00144239 |
| Fa2h | 7,863464422 | 0,729353204 | 0,00000519 | 0,001869891 |
| Cldn11 | 9,659859378 | 0,804930069 | 0,00000541 | 0,001910377 |
| Pde10a | 10,19689363 | 0,396871906 | 0,00000609 | 0,002055066 |
| Ntf3 | 4,744423666 | -0,769804677 | 0,0000074 | 0,002359843 |
| Cnksr3 | 7,061357185 | 0,581960942 | 0,00000789 | 0,002467867 |
| Gsn | 9,496893965 | 0,657798115 | 0,00000835 | 0,002564693 |
| Erbb3 | 6,63106478 | 0,638710738 | 0,00000892 | 0,002639535 |
| Thbs4 | 4,459510775 | 0,949831152 | 0,00001 | 0,002813664 |
| 1700016P03Rik | 6,371516477 | -0,596772857 | 0,0000112 | 0,003089316 |
| Nkx6-2 | 4,930460027 | 0,915398433 | 0,0000142 | 0,003810841 |
| 1810049J17Rik | 5,285190416 | -0,858678538 | 0,0000165 | 0,004270157 |
| Ddc | 5,586004781 | 0,734670344 | 0,0000173 | 0,004350241 |
| Cnp | 11,27643462 | 0,618838219 | 0,0000188 | 0,004645962 |
| Gm4409 | 5,519508287 | -0,561333921 | 0,0000192 | 0,004693585 |
| Hipk4 | 5,24654989 | -0,544687464 | 0,0000233 | 0,005590229 |
| Rnf122 | 7,312323235 | 0,544814982 | 0,0000241 | 0,005713819 |
| Fbxo32 | 6,923939992 | 0,560622746 | 0,0000248 | 0,005781423 |
| Rasgef1b | 9,64835912 | 0,410577466 | 0,000028 | 0,006371634 |
| Mbp | 13,91411933 | 0,777938255 | 0,0000299 | 0,006711352 |
| Adora2a | 6,022648217 | 0,584921755 | 0,0000367 | 0,008088682 |
| Elovl7 | 6,926672412 | 0,620958585 | 0,0000371 | 0,008088682 |
| Slc45a3 | 3,781312709 | 1,000140868 | 0,0000411 | 0,008743406 |
| 1700047M11Rik | 5,468802989 | 0,833857795 | 0,0000434 | 0,009119293 |
| Gm15337 | 4,768984581 | -0,682221596 | 0,0000452 | 0,009363815 |
| Tpx2 | 5,948555967 | 0,609754926 | 0,0000526 | 0,010621885 |
| Mal | 8,407818278 | 0,903169585 | 0,0000519 | 0,010621885 |
| 4930404N11Rik | 5,227762848 | -0,734521179 | 0,0000583 | 0,011512271 |
| Erbb2ip | 9,421443092 | 0,409296601 | 0,0000672 | 0,012804779 |
| Anln | 5,864631814 | 0,674209603 | 0,0000999 | 0,018411831 |
| Plekhh1 | 7,251100575 | 0,491337231 | 0,0000997 | 0,018411831 |
| Mob3b | 6,265102048 | 0,640626217 | 0,000105023 | 0,018932366 |
| Tmem88b | 8,052224336 | 0,695812026 | 0,000105028 | 0,018932366 |

| | | | | |
|---|---|---|---|---|
| **Sgk2** | 3,210154152 | 1,328355184 | 0,000117092 | 0,020880206 |
| **Carhsp1** | 7,645317422 | 0,535047937 | 0,000144918 | 0,025567205 |
| **Kif19a** | 6,994888974 | 0,549714314 | 0,000166517 | 0,029068544 |
| **Syndig1l** | 6,65459875 | 0,517832345 | 0,000174901 | 0,029902589 |
| **Casr** | 4,988691236 | 0,711601482 | 0,000207125 | 0,03302854 |
| **Cyb5d1** | 7,030386252 | 0,483482965 | 0,00021898 | 0,034586384 |
| **RP23-332E2.7** | 3,729200235 | -0,923109694 | 0,00022695 | 0,035506921 |
| **Oprk1** | 5,600679909 | 0,655892667 | 0,000239432 | 0,037109723 |
| **Mir212** | 3,357725456 | -1,357881457 | 0,00025662 | 0,038892967 |
| **Lox** | 5,30762312 | -0,534184372 | 0,000257973 | 0,038892967 |
| **Tnni1** | 3,584368874 | 0,923998415 | 0,000288401 | 0,041954719 |
| **Rgs9** | 8,038663943 | 0,44796957 | 0,000305048 | 0,043766787 |
| **Gm4737** | 3,95653166 | -0,788699017 | 0,000316984 | 0,044843778 |
| **RP24-390L20.5** | 3,288289841 | -0,897992141 | 0,000322475 | 0,044940499 |
| **Gm10475** | 4,581419909 | -0,77032687 | 0,000330416 | 0,045663486 |
| **Gm10143** | 4,741120126 | -0,632053476 | 0,000337194 | 0,046151657 |
| **Ppp1r14a** | 4,267036226 | 0,85165977 | 0,000351585 | 0,047403969 |
| **Depdc1b** | 3,045036993 | 1,258004209 | 0,000362975 | 0,048156581 |
| **Bcas1** | 10,9752945 | 0,446732673 | 0,000361205 | 0,048156581 |
| **Gm9887** | 3,284525342 | -1,036015739 | 0,000379917 | 0,049574412 |
| **Slc6a20b** | 3,805006358 | -0,896122701 | 0,000382629 | 0,049574412 |
| **Ube4bos3** | 4,06282201 | -0,737740914 | 0,000377679 | 0,049574412 |

### Supplementary Table 2: Primers used for RT-qPCR

| Gene Name | Forward | Reverse |
|---|---|---|
| **Arc** | CTGAAGCAGCAGACCTGACA | CTCAGCAGCCTTGAGACCTG |
| **Plp1** | GCAAAGTCAGCCGCAAAACA | GCCCCTACCAGACATCTAGC |
| **Mag** | TTCTCAGGGGGAGACAACC | ACTCTCCTGGGGCTCTCAGT |
| **Mog** | CTGGCAGGACAGTTTCTTGA | AAAGAGGCCAATGGGAAATC |
| **Fos** | TACTACCATTCCCCAGCCGA | GCTGTCACCGTGGGGATAAA |
| **Fosb** | CTTCAACCAGCACAACCACC | TCTGCGAACCCTTCGCTTTT |
| **Gapdh** | CTTCTTGTGCAGTGCCAGC | GAGGTCAATGAAGGGGTCGT |
